# High distinctness of circadian rhythm is related to negative emotionality and enhanced neural processing of punishment-related information in men

**DOI:** 10.1101/2024.09.13.612861

**Authors:** Patrycja Scislewska, Michal R. Zareba, Julia Lengier, Aaron E. Schirmer, Piotr Bebas, Iwona Szatkowska

## Abstract

For years, research on the human biological clock has focused primarily on chronotype (phase of the circadian rhythm). However, a second dimension – distinctness (subjective amplitude) of the rhythm, has so far been overlooked. This study aimed to explore the intricate interplay between psychometric traits and reward-punishment processing, considering both chronotype and distinctness. Circadian rhythmicity characteristics of 37 healthy men (aged 20-30) were measured using the Morningness-Eveningness-Stability-Scale improved (MESSi) questionnaire. We also employed a battery of psychometric questionnaires and used functional magnetic resonance (fMRI) during the Monetary Incentive Delay task, which is a common method of assessing reward-punishment processing. We found a positive association between distinctness and the activity in the bilateral Superior Frontal Gyrus (SFG), Supplementary Motor Area (SMA) and Ventral Tegmental Area (VTA) during processing of punishment cues. These results are consistent with psychometric findings – a significant positive association between distinctness and sensitivity to punishment, neuroticism, behavioral inhibition system (BIS), attention deficits, and negative emotionality. As regards eveningness, we found its negative association only with sensitivity to punishment and BIS. These results highlight the crucial role of distinctness in human functioning, especially in terms of punishment processing and negative emotionality.

## Introduction

Circadian rhythmicity in humans is represented by a complex phenotype with varying levels of alertness, mood and motivation observed across a 24 hour cycle. To describe these daily rhythms a characterization of at least two separate dimensions: chronotype (rhythm phase) and distinctness (subjective amplitude of the rhythm) would be needed^1^. The chronotype represents an individual’s preferred time of activity (i.e., morningness-eveningness), while the distinctness – the strength of this preference, understood as the subjective amplitude between hyper- and hypo-activation phases (e.g., high distinctness will manifest itself in the feeling that “There are moments during the day when I feel unable to do anything”, while low distinctness – “I can focus at any time of the day”)^2–4^.

Although both components can influence behavior, to date, the vast majority of investigations have solely used a unidimensional assessment of morningness-eveningness^5^, while distinctness has remained overlooked. In those studies, eveningness was shown to be associated with negative moods and affect^6,7^, increased anxiety^8^, propensity to depression^8–10^, more use of substances, including alcohol, nicotine, and caffeine^8,11^, and vulnerability to addiction^12^. Propensity to negative emotionality, depression and addiction might be associated with deficits in reinforcement processing and altered reward-related brain function that were observed in people with evening chronotype^13^.

While there has been significant research pointing to the associations between morningness-eveningness, negative emotionality and deficits in reinforcement processing, investigations dealing with distinctness are rather sparse. However, recent research has emphasized the role of distinctness, which correlates negatively with life satisfaction and positively with negative emotionality^14–16^. Moreover, higher subjective amplitude values are positively correlated with past negative time perspective and negatively correlated with focusing on the future^17^. Relationships between distinctness and negative emotionality, more spontaneous mind wandering, and neuroticism have also been observed^18–21^. Importantly, higher distinctness may be more strongly related to negative emotionality than eveningness preference^14^. Although such results point to the relationship between the level of distinctness and efficiency of reinforcement processing, this relationship has never been directly tested.

To better understand the complex nature of human circadian rhythmicity, the aims of this study were to: (1) concurrently test associations between the circadian rhythm components of eveningness and distinctness, negative emotionality, and alterations in reinforcement processing, and (2) explore the neural basis of observed circadian-related differences in the reinforcement processing. The investigation consisted of two parts – examining the subjects’ behavior using psychometric questionnaires and measuring brain activity using functional Magnetic Resonance Imaging (fMRI). Building on the recent findings, it was expected that variability in affective processing would be more strongly associated with distinctness than with eveningness. To verify this hypothesis, we employed a battery of psychometric questionnaires, which allowed us to assess individuals’ sensitivity to punishment and reward, personality traits, positive and negative affect, behavioral inhibition system and behavioral activation system (BIS/BAS), anxiety levels, and attention deficits (for details, see: Methods). To determine neural correlates we decided to use fMRI during the Monetary Incentive Delay (MID) task (Fig. 1.), which is a common method of assessing reward- and punishment-motivated behavior^22,23^. During the task performance, participants had to rely on previously learned associations between the cues and monetary outcomes, therefore, the task is assumed to activate brain regions involved in reward-punishment processing, as well as attention and working memory, such as prefrontal cortex (PFC). Moreover, accumulating evidence suggests the part of the mesolimbic pathway – the ventral tegmental area (VTA) – is not only a hub for reward regulation^24^, but also serves as a circadian oscillator that regulates reward and motivation across the day^25^, thus we expected to find differences in VTA activity during reinforcement anticipation phase. To investigate our hypotheses, we used both whole-brain and region-of-interest (ROI) analysis approaches.

**Figure 1.**
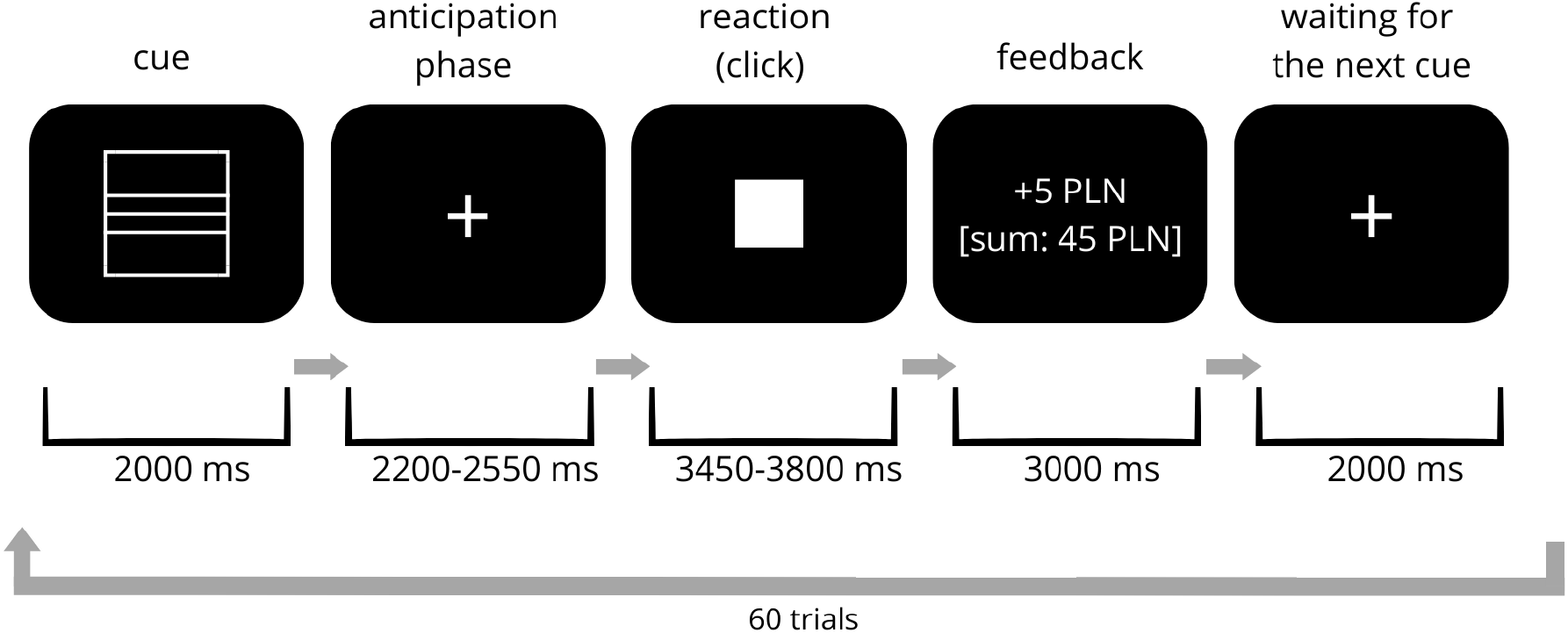
The Monetary Incentive Delay task paradigm. At the beginning of each trial, participants obtain the graphical cue with assigned monetary value – reward cues (square with 3 lines represents big win: +5 PLN (Polish Zloty, currency in Poland, 1 PLN equals 0.30 USD), square with 1 line – small win: +1 PLN), punishment cues (circle with 3 lines represents big loss: –5 PLN, circle with 1 line – small loss: –1 PLN). After the anticipation phase, the white-filled square on the scanner screen is displayed, signaling the moment of reaction – the participant should click on the pad as fast as possible to gain a reward or avoid punishment, depending on the monetary cue. There are also oddball trials (+/–10 PLN) in which inhibition of the automatic reaction is required. The oddball trials were not analyzed in terms of brain activity due to their low repetition number, and their main goal was to keep participants focused on correctly processing the meaning of cues. The task consisted of 60 trials. For details, see: Methods.

## Results

The subjective characteristics of the participants’ circadian rhythms were assessed using the subscales for eveningness and distinctness. The higher the score in the eveningness subscale, the more evening-oriented the individual is expected to be. Similarly, the higher the score in the distinctness subscale, the stronger the within-day fluctuations in mental processes. Additionally, the sleep quality of the participants was also controlled. The values were distributed normally according to the Kolmogorov-Smirnov test (p > 0.05). The demographic summary of the sample is presented in Table 1.

**Table 1.** Demographic summary of the cohort (37 male participants) used in the study. Eveningness and distinctness were measured using the Morningness-Eveningness-Stability-Scale improved (MESSi) questionnaire^4,26^. Theoretical ranges of eveningness and distinctness subscales were 5-25. Sleep quality was controlled using Athens Insomnia Scale (AIS)^27^ with theoretical range 0-24. Abbreviation: SD - standard deviation.

| Variable (n = 37) | Value range | Mean ± SD | Median |
| --- | --- | --- | --- |
| Age (years) | 20-30 | 24.4 ± 2.1 | 25 |
| Eveningness | 10-24 | 18.6 ± 4.6 | 20 |
| Distinctness | 10-25 | 18.2 ± 3.0 | 18 |
| AIS | 1-16 | 5.6 ± 3.2 | 5 |

### Behavioral results

To assess the effect of distinctness and eveningness on these psychometric parameters, an Ordinary Least Squares (OLS) regression analysis across multiple dependent variables was performed. Distinctness has a statistically significant positive effect on the following dependent variables: sensitivity to punishment, neuroticism, openness, BIS, and attention deficits, while eveningness has a statistically significant negative effect on sensitivity to punishment and BIS. Coefficient values and confidence intervals from the OLS regression analysis for all of the measured psychometric dependent variables are shown in Fig. 2. Detailed data for all psychometric values are in Supplementary Table S1. These results support the hypothesis that distinctness may act as a more important factor than eveningness in context of affective processing, as more parameters varied in distinctness when compared to eveningness.

**Figure 2.**
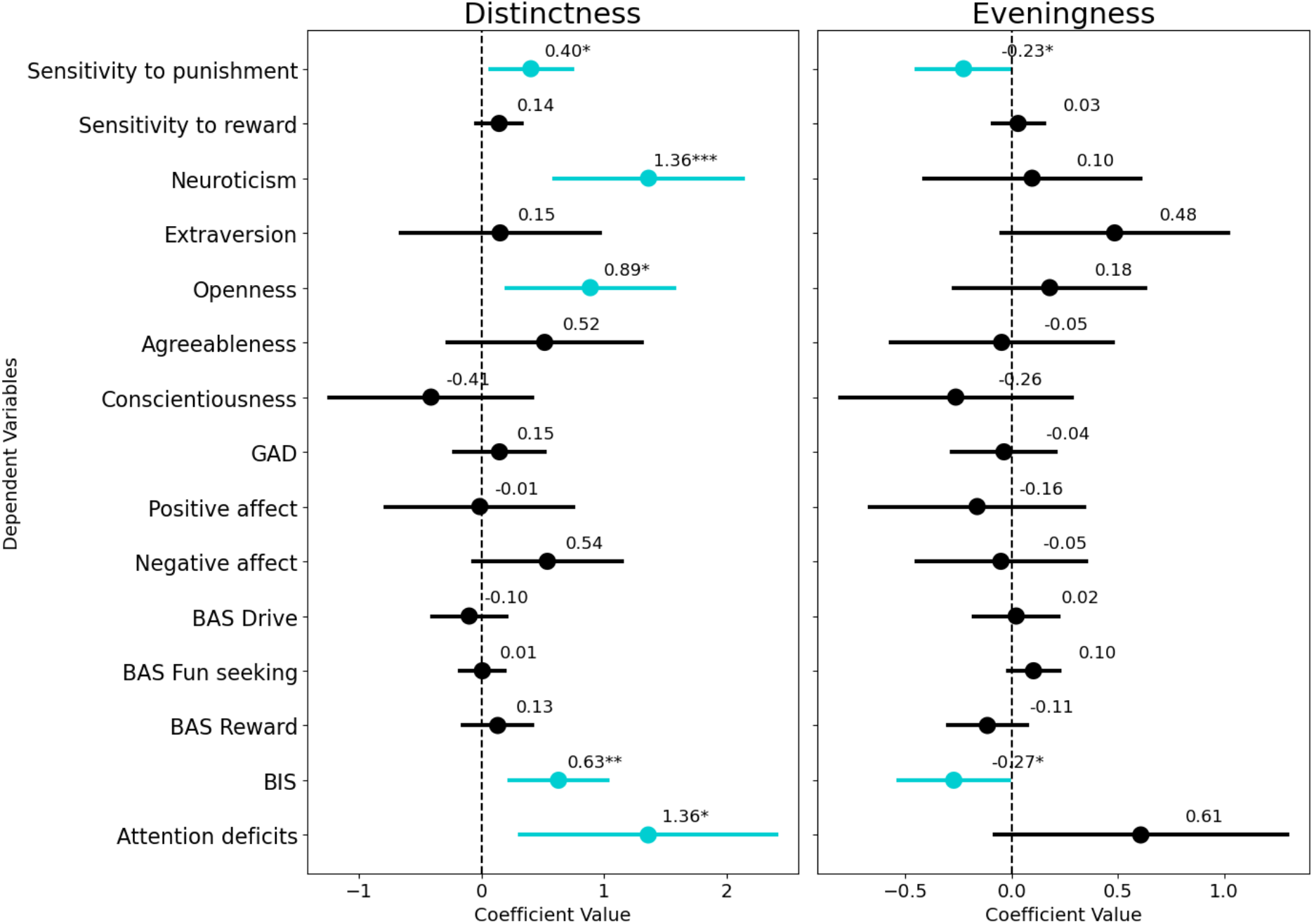
The Ordinary Least Squares (OLS) regression results for all psychometric parameters. Distinctness (left panel) and eveningness (right panel) were used as predictor variables. Dots and numbers represent coefficient values, horizontal bars - confidence intervals (95% CI). Statistically significant associations are marked in blue, * represents *p*-value ≤ 0.05, ** represents *p*-value ≤ 0.005, *** represents *p*-value ≤ 0.001. Psychometric parameters were assessed using the Sensitivity to Punishment and Sensitivity to Reward Questionnaire (SPSRQ^28,29^), NEO-Five Factor Inventory (NEO-FFI^30,31^), Generalized Anxiety Disorder (GAD-7^32^), Positive and Negative Affect Schedule (PANAS^33,34^) Behavioral Inhibition System and Behavioral Activation System (BIS/BAS^35,36^), and Adult ADHD Self-Report Scale (ASRS^37,38^).

According to previous studies, distinctness usually appeared in aspects of negative emotionality, and not positive ones. Our OLS analysis results motivated us to investigate this phenomenon. To confirm this distinction, we selected pairs of psychometric properties which have corresponding subscales for negative and positive emotionality – neuroticism / extraversion, BIS / BAS, negative affect / positive affect, and compared them across different circadian features.

We divided participants into groups of higher / lower distinctness and higher / lower eveningness using a median split and conducted a 2 {distinctness groups} × 2 {eveningness groups} ANOVA. The two way ANOVA demonstrated a statistically significant direct effect of distinctness on neuroticism (*F*(1,33) = 8.71, *p* = 0.006), BIS (*F*(1,33) = 6.16, *p* = 0.018), and negative affect (*F*(1,33) = 4.266, *p* = 0.047). No significant associations were found between distinctness and any of the positive emotionality subscales (Fig. 3.), as well as between eveningness and any psychometric traits. There were no interactions between distinctness and eveningness. Detailed data are presented in Supplementary Table S2. These results underline the complex interplay between circadian rhythmicity and human emotionality and highlight the diverse associations with its negative and positive facets.

**Figure 3.**
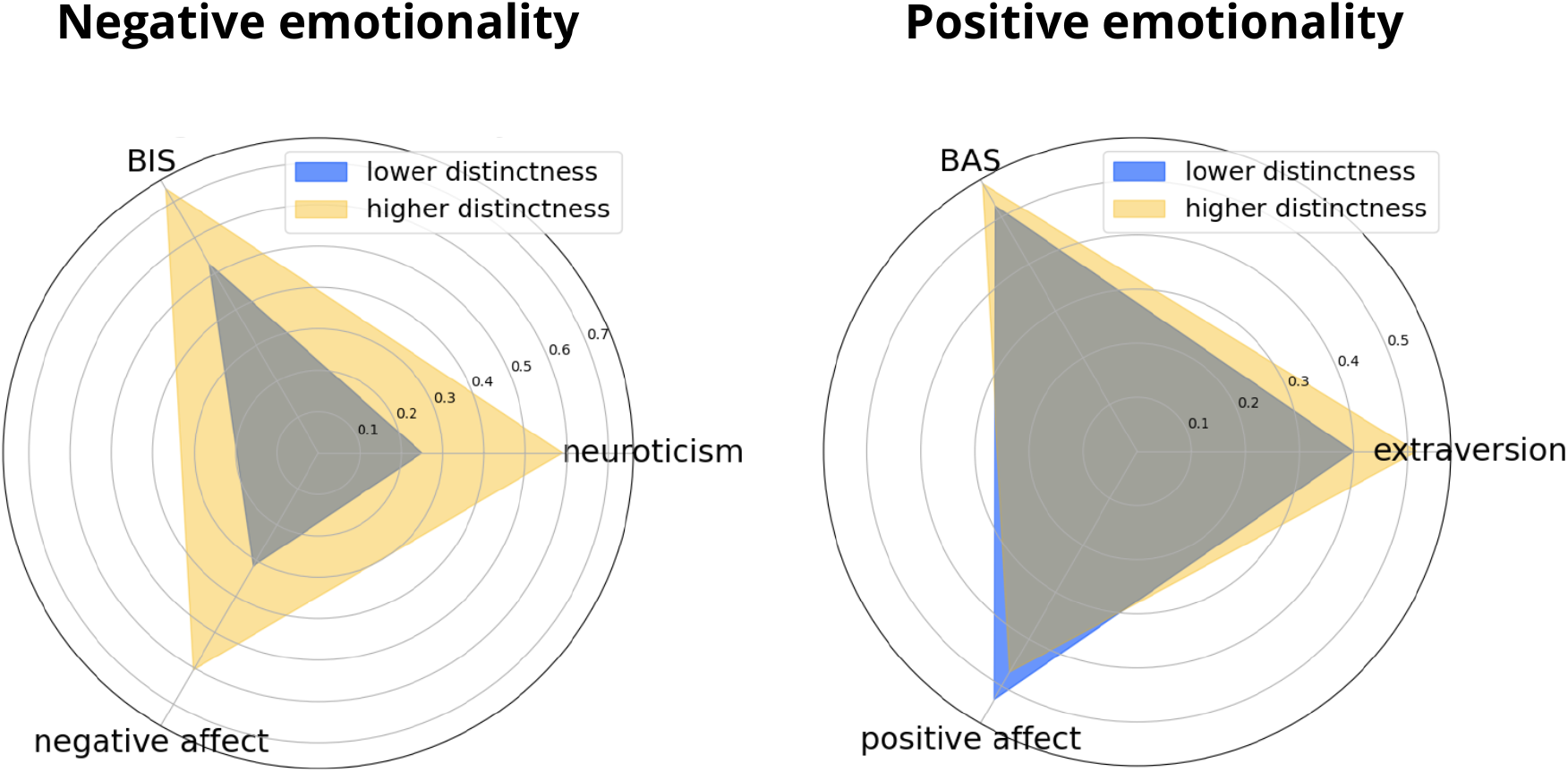
Differences in negative (left panel) and positive (right panel) emotionality between lower-distinctness (blue) and higher-distinctness (yellow) individuals. Two groups of distinctness values were obtained using median split. Psychometric parameters values shown in the corner are normalized medians of these parameters in each group. All of the negative aspects of emotionality differ significantly between groups of higher and lower distinctness. No statistically significant difference was found in case of positive emotionality.

### Neuroimaging results

Considering the associations observed between circadian characteristics, sensitivity to punishment, and negative emotionality, we expected to find neuronal differences in affective processing. In order to determine neural correlates of these explored processes, a Monetary Incentive Delay (MID) task was employed (for details, see: Methods). We focused on anticipation phases of the MID task and defined the same contrasts as was done in previous studies^22,23^, i.e. we contrasted (1) each trial type (reward / punishment) against the implicit baseline both individually for each value and collapsed across values of the trial type, (2) high value trials against low value trials within each trial type, (3) reward anticipation trials against loss anticipation trials. The feedback phase of the MID task was not a subject of interest in the current study.

To investigate our hypotheses, we used both whole-brain and region-of-interest (ROI) analysis approaches. To mitigate factors that may potentially modulate the functionality of the investigated brain regions (e.g., the synchrony effect^39^ or seasonal changes in mood and cognitive performance^40^), all fMRI examinations were carried out from 1 PM to 5 PM during two weeks in summer (for details, see: Methods).

The whole-brain neuroimaging results revealed associations between distinctness and negative reinforcement processing, while controlling for eveningness. Distinctness was positively associated with the activity of the bilateral superior frontal gyrus (SFG) and supplementary motor area (SMA) during the anticipation phase of the punishment trials (across all trial values), Fig. 4a. The whole-brain analysis results suggest the involvement of brain areas related to working memory and attention^41,42^, which is consistent with the predictions – to effectively perform the task, participants were required to remember the meaning of monetary cues. However, this approach did not reveal differences in activity of the brain regions involved in the reward-punishment processing. In each trial, participants experienced a reward or punishment anticipation phase, therefore, the region-of-interest (ROI) analysis was performed, in which we explored the activity of the ventral tegmental area (VTA). An Ordinary Least Squares (OLS) analysis performed with distinctness and eveningness as predictors using the ROI approach showed the distinctness-related difference in the VTA activity during the anticipation of loss vs. anticipation of win trials, Fig. 4b. There were no significant neuroimaging findings associated with eveningness. The results are detailed in Table 2. and Fig. 4.

**Table 2.** Significant associations between the distinctness and neuroimaging results. Abbreviations: B, bilateral. ^a^Whole-brain analysis corrected for multiple comparisons with cluster-level family-wise error rate approach. ^b^Region-of-interest analysis.

| Contrast | MNI (x, y, z) | Area | Voxels | Statistics | p-value |
| --- | --- | --- | --- | --- | --- |
| Loss anticipation > implicit baseline | 0, 24, 62 | B superior frontal gyrus (SFG)<br>B supplementary motor area (SMA) | 261 | Z = 3.44 | 0.033<sup>a</sup> |
| Loss anticipation > win anticipation | 0, -17, -14 | B ventral tegmental area (VTA) | 32 | T = 3.03 | 0.005<sup>b</sup> |

**Figure 4.**
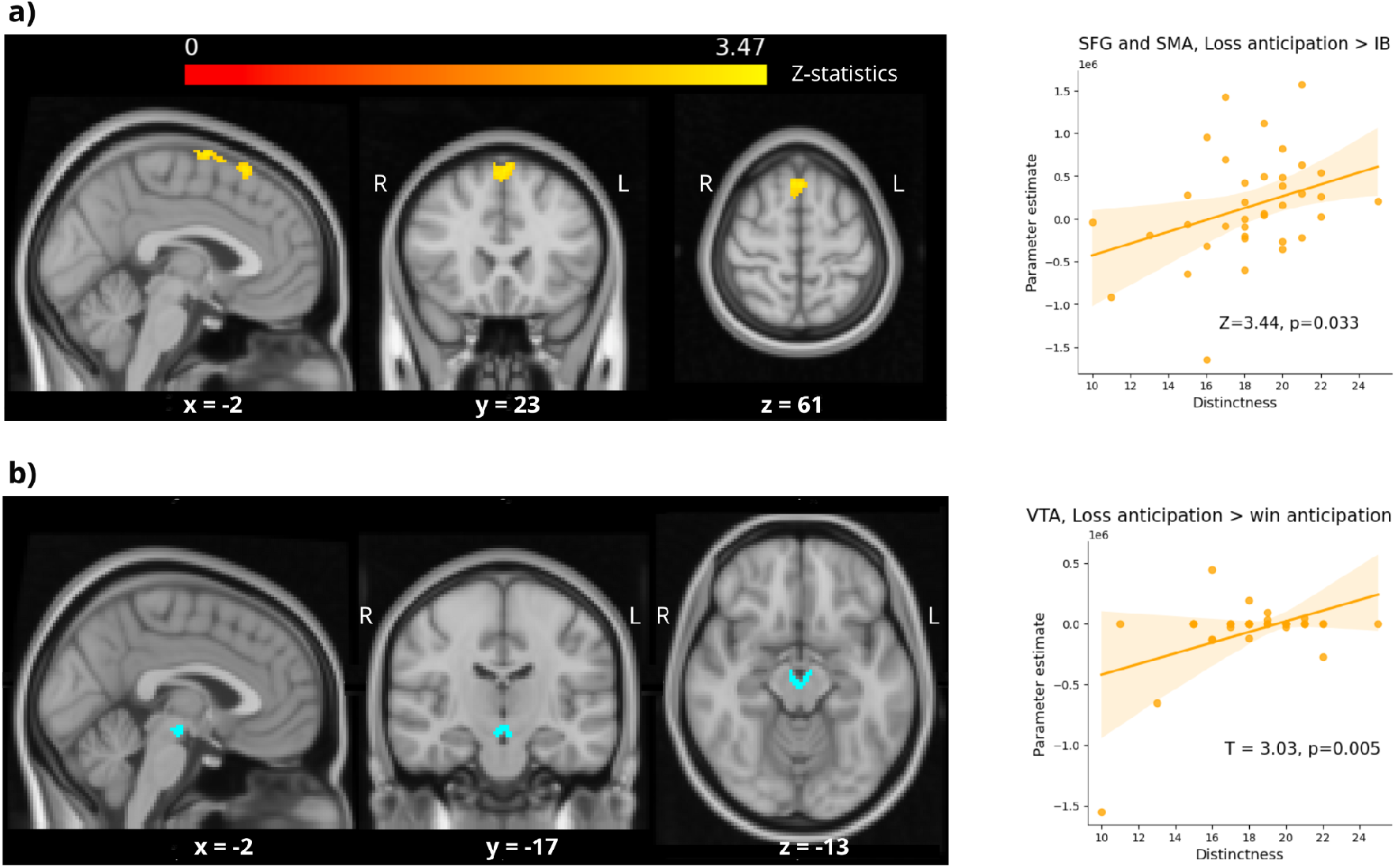
Neuroimaging results. Top panel (a): bilateral superior frontal gyrus (SFG) and supplementary motor area (SMA) activity in anticipation of loss phase is positively associated with the distinctness of the circadian rhythm. Bottom panel (b): differences in the activity between the anticipation of loss vs. anticipation of win trials for the ventral tegmental area (VTA) are positively related to distinctness. Activations are shown on standard human brain template (MNI152). Coordinates (x, y, z) were chosen to show the best representation. Linear regression lines are shown on the charts.

In our dataset, 3 participants obtained Athens Insomnia Scale values higher than 10, which indicated potential chronic sleep deprivation. To check whether their brain activity was significantly impacted, we re-conducted the whole brain and region-of-interest analyses excluding these participants. The results remained unchanged (see: Supplementary Table S3).

All of the relationships found between distinctness and brain activity occurred during the anticipation of loss phase, which aligns with previously described behavioral observations of elevated scores in sensitivity to punishment and negative emotionality of individuals characterized by high distinctness values. Moreover, in the behavioral part of this study, we showed that attention deficits were positively related to the distinctness of the circadian rhythm, which may explain the increased activity in SFG (Superior Frontal Gyrus) and SMA (Supplementary Motor Area) – brain areas crucial for attention and working memory during task performance^41,42^.

### Monetary Incentive Delay (MID) task behavioral results

No statistically significant relationships were found between distinctness or eveningness scores and (I) final monetary score, (II) number of incorrect responses (omitting responses when due to provide one, and failure to suppress responses when indicated to do so), (III) average reaction times across all anticipation trials, (IV) average reaction times in any conditions of reward or punishment anticipation phase using two-way ANOVA. Detailed information is included in the Supplementary Table S4.

## Discussion

This study reveals that individual differences in circadian rhythm distinctness are reflected in neural activity during the performance of a task activating affective processes, which sheds new light on the neural correlates of circadian rhythm variability. Namely, higher levels of distinctness are related to preferential processing of punishment cues in the dopaminergic midbrain (Ventral Tegmental Area, VTA) and two parts of frontal cortex (Superior Frontal Gyrus, SFG, and Supplementary Motor Area, SMA). This characteristic is supported by positive associations between the subjective amplitude of the rhythm with neurotic personality type and sensitivity to punishment, elevated scores on the behavioral inhibition system, as well as increased negative affect (Fig. 2. and Fig. 3.). Therefore, we confirm the previously established relationship of high subjective amplitude with negative emotionality^18–21^, and provide evidence for the association between a higher value of distinctness and enhanced neural processing of punishment-related information. Additionally, we extend the state of knowledge by reporting the association of distinctness with openness to experience and attention deficits, two other cognitive domains that are also tied to dopaminergic functioning^43,44^.

Our findings of preferential processing of negatively-valenced stimuli in participants with higher distinctness are likely related to increased activity of the salience-encoding subsystem in response to the punishment cues, i.e. they are perceived as motivationally more important. According to loss aversion theory, individuals prefer to avoid loss rather than acquire a gain and it is even suggested that losses are psychologically twice as harmful as gains are beneficial^45^. In light of this theory, our results might suggest that people with higher distinctness manifest higher levels of loss aversion. Loss avoidance plays an important role in mental disorders such as ADHD^46^, which in our study might be reflected in positive relation between distinctness and attention deficits measured using Adult ADHD Self-Report Scale. Loss aversion may also affect working memory performance through the amplification of activity within prefrontal and visual association regions selective to processing the stimuli to be remembered^47^. As the anticipation phase of the MID task contains working memory components, our results might reflect the modulation of working memory functions by motivation through loss aversion in individuals with higher distinctness. This corresponds well with the functions played by frontal regions associated with distinctness in our study. The SFG and SMA are involved in executive functions^48^ and in preparation for the adequate motor response to avoid punishment^49^. In the broader context, however, these brain areas are additionally involved in emotion regulation^50^. Thus, the increased neural reactivity to negative stimuli might in the long run contribute to maladaptive emotion processing, and as such predispose individuals with greater distinctness to developing affective disorders^6^.

The particular role of distinctness in the processing of negative emotions, rather than emotionality in general, is emphasized by the fact that not a single association was found between distinctness and broadly understood positive emotionality (Fig. 3). This characteristic was reflected in the brain imaging results, where distinctness was positively linked to the activity of frontal cortices and VTA during the anticipation phase of loss trials, despite the lack of any associations with behavioral task measures. Combined with the questionnaire data, these findings indicate that the increased neural resources dedicated to the processing of punishment cues in individuals with high distinctness might not be necessary for adequate task performance but may rather reflect a bias towards the processing of negatively-valenced stimuli. These results are complementary to our previous work^51^, in which we showed negative correlation between subjective amplitude and both grey matter volume and cortical thickness in the left primary visual cortex, which plays a crucial role not only in emotional perception but also in the down-regulation of negative emotions^52^.

Currently, the exact mechanism through which the amplitude of diurnal variations in mood and cognition could contribute to increased dopaminergic processing of negative stimuli remains unknown. Midbrain dopamine system, especially the ventral tegmental area (VTA), contains separate circuitries encoding reinforcement value and motivational salience, forming the circuit basis of adaptive and maladaptive motivated behaviors^53^. As greater sensitivity to light is linked to more pronounced fluctuations in the levels of diurnally secreted hormones^54^, the non-image forming visual system is a perfect candidate to mediate the discussed effects. In diurnal rodents, the influence of light on depression- and anxiety-like behaviors is translated through orexinergic neurotransmission^55^, and orexins increase both the activity of the dopaminergic neurons and the amplitude of molecular circadian rhythmicity in habenula^56,57^, providing multiple pathways for potential modulation of affective processing.

While the proposed associations between light and distinctness of the circadian rhythm remains speculative, it corresponds well with the findings regarding light therapy in humans with seasonal affective disorder (SAD) and depression. Light therapy is the treatment of choice for SAD, however, some people, such as evening-oriented individuals with larger amplitude of diurnal mood variation, benefit from the treatment more than others^58^. The variability in the dopaminergic system may explain the winter-related decrease of positive emotions and attention^59^, as seasonal fluctuation of striatal dopamine synthesis and D2R availability have been suggested as a major factor in the development of SAD^60^. These findings suggest that photoperiod changes, particularly variability in the spectral composition of natural light^61^ and total light irradiance, can fundamentally alter monoaminergic systems in the brain, leading to changes in mood-related behavior^62^.

Investigations linking circadian rhythmicity with negative mental health outcomes have traditionally focused on the role of eveningness preference^12,13,18,63^. Nevertheless, there is a growing evidence that the effects associated with late circadian phase are to a great extent mediated by sleep quality^64,65^ and their magnitude is lower compared to the influence of distinctness^6,16^. In the current sample, neither the circadian time preference (eveningness) nor subjective amplitude (distinctness) were related to sleep complaints, which might be related to the inclusion of only male participants^66,67^. The current study, as the first task-based fMRI study, focused solely on male participants to reduce the number of factors that may influence affective processes, e.g., women’s hormonal cycles^68^). However, further studies should include female individuals, as Smarr and collaborators showed that women are not more behaviorally variable than men, thus ovarian cyclicity is not the driver of chronotype-related sex variability^69^. Moreover, in the current study, all the data were acquired during the summer, so further winter data collection and longitudinal research are required.

## Conclusions

The current study indicates that distinctness (subjective amplitude) of the rhythms may be more tightly related to the psychometric measures of behavior than eveningness (rhythm phase), which has been the primary focus of research so far. It extends the state of knowledge by showing that increased neural processing of punishment cues depends on distinctness. This finding is consistent with the previously reported links of higher rhythm amplitude with negative emotionality, which we also confirmed in the current study. Moreover, we demonstrated that distinctness of the circadian rhythm is linked to openness to new experiences and attentional deficits. As such, the current study highlights the complex, multidirectional, and non-mechanistic nature of the relationships between circadian rhythmicity and emotionality. By integrating distinctness as an additional factor into analyses, we can achieve a more nuanced and comprehensive understanding of human circadian behavior, potentially uncovering new insights and opportunities for targeted interventions aimed at improving behavioral outcomes and quality of life.

## Methods

### Participants

The described study involved 37 healthy men. All subjects were aged 20-30, had normal or corrected-to-normal vision, were drug-free, and self-reported no history of either neurological, nor psychiatric disorders. The described protocol was approved by Rector’s Committee for the Ethics of Research Involving Human Participants at the University of Warsaw (decision no. 230/2023) and was performed in accordance with the Declaration of Helsinki (World Medical Association, 2008). Written informed consent prior to experimentation was obtained from all participants.

### Circadian characteristics

The subjective characteristics of the circadian rhythms of the participants were assessed using the Polish version of the Morningness-Eveningness-Stability-Scale improved (MESSi) questionnaire^4,26^, which includes subscales for Eveningness (rhythm phase), Distinctness (rhythm amplitude) and Morning Affect. However, in this study we focused on an in-depth analysis of the role of eveningness and distinctness of the circadian rhythm, following Carciofo’s findings^6^, according to which distinctness and morning affect can be treated as separate predictors. Distinctness measures the subjective amplitude or the range of diurnal variation. This variation results from the ability to volitionally modulate one’s own psychophysiological state and feel the difference between hyper- and hypo-activation phases (e.g., “I can focus at any time of the day”; “There are moments during the day where I feel unable to do anything”). Higher scores show higher fluctuations. Each dimension was tested with 5 objects ranked from 1 to 5, which are summed together, making the theoretical score range from 5 to 25 points. The higher the score in the eveningness subscale, the more evening-oriented the individual is. Similarly, the higher the score in the distinctness subscale, the stronger the within-day fluctuations in mental processes. In the current dataset, the eveningness scores ranged from 10 to 24, while distinctness scores varied from 10 to 25. Additionally, the sleep quality of the participants was controlled using the Athens Insomnia Scale (AIS) questionnaire^27^. In our cohort, the score range was 1 to 16, while the theoretical range is 0 to 24.

### Behavioral data

The individual differences in reinforcement processing were assessed using the Sensitivity to Punishment and Sensitivity to Reward Questionnaire (SPSRQ^28,29^). Additionally, to better characterize the link between the emotional processing and circadian functioning, we obtained complementary data with the use of: NEO-Five Factor Inventory (NEO-FFI^30,31^), Behavioral Inhibition System and Behavioral Activation System (BIS/BAS^35,36^), Positive and Negative Affect Schedule (PANAS^33,34^), Generalized Anxiety Disorder (GAD-7^32^) and Adult ADHD Self-Report Scale (ASRS^37,38^).

### Monetary Incentive Delay (MID) task

Each trial in the MID task consisted of 5 stages as shown in Fig. 1 (in the Introduction section). At the beginning of each trial, a graphic cue indicative of its monetary value was displayed in the scanner.

During the trial, participants had to rely on previously learned associations between the cues and monetary outcomes. Following the cue, the anticipation phase occurred, which consisted of a fixation point (represented as a “+” in Fig. 1.) displayed in the center of the screen. The duration of the anticipation was pseudorandomized to avoid subjects learning the clicking rhythm. In the next stage (“reaction (click)” in Fig. 1), a screen with a white, filled square appeared in the place of the fixation cross, and the subjects were instructed to press the response button as quickly as possible to obtain a reward or avoid punishment (depending on the previously displayed cue). After their reaction, participants received the feedback, displaying the result of a given trial and the total amount won so far. After the feedback phase, a fixation point (represented as a “+” in the panel entitled “waiting for the next cue” in Fig. 1). appeared to mark the end of the trial. The complete duration of each trial was 13 seconds.

The task consisted of 60 trials: 28 trials with reward cues (large win: +5 PLN, small win: +1 PLN), 28 trials with punishment cues (large loss: –5 PLN, small loss: –1 PLN) and 4 oddball trials (+/–10 PLN). The oddball trials were not analysed in terms of the brain activity due to their low repetition number, and their main goal was to keep participants focused on correctly processing the meaning of cues. The sequence in which the cues were presented in the task was pseudo-randomized, but consistent across all subjects (the task code utilised a predefined list dictating the order of cue presentation). A summary of these cues is illustrated in Fig. 5. Participants were shown an identical screen (translated into Polish) at the commencement of the MID task session.

**Figure 5.**
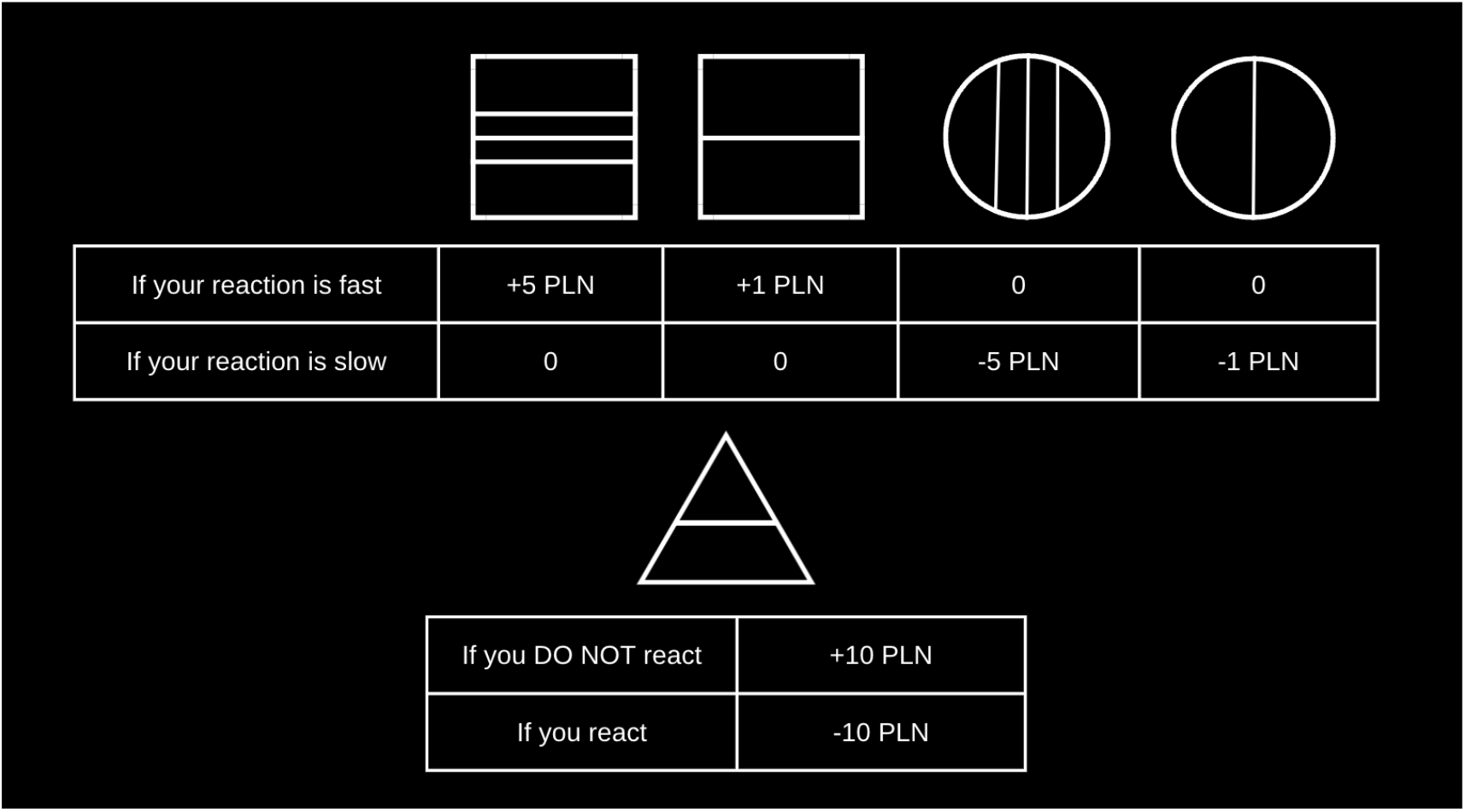
Description of the cues from the Monetary Incentive Delay (MID) task. Participants were taught the meaning of these graphical cues during a training session. At the beginning of the neuroimaging session, participants were shown this exact screen to give them the opportunity to recall the monetary value of the cue. However, during the MID task, participants had to rely on their memory.

Prior to entering the scanner, the participants became thoroughly familiar with the task by memorizing the values assigned to specific graphical symbols (cues), and becoming accustomed to providing answers in the same manner as during the subsequent scanning session (the index finger of the right hand). The training session additionally enabled the measurements of the individual reaction times, which were used to tailor the threshold in the MID task to achieve a 60% success rate in the scanner for each participant. This approach was meant to ensure that the participants would be able to win in a sufficient number of trials for the reinforcement processes to take place, simultaneously demanding their continuous focus on the task. This goal was additionally ensured by informing participants that their financial remuneration for participation in the study would be equal to the amount of total gain in the MID task. Participants were told that the guaranteed reward for participating in the study was 50 PLN (13 USD), and that the total final payout they would receive would depend on their performance in the experiment (up to 110 PLN, 30 USD).

### Data acquisition

The MRI data was acquired using 3T MAGNETOM Prisma Siemens scanner at the Laboratory of Brain Imaging (LOBI), Neurobiology Center at the Nencki Institute of Experimental Biology Polish Academy of Sciences.

Scans were performed using a 32-channel head coil. The subjects held a response pad in their right hand. The right button of the response pad was used to complete the task. To ensure optimal visual acuity, participants were offered MRI-compatible glasses if necessary. For each subject T1-weighted anatomical scan, fieldmap, and T2*-weighted scans during tasks were performed. T1-weighted anatomical scans were performed using Echo Planar Sequence (GR\\IR), Slice Thickness = 1 mm, TE = 0.00296 s, TR = 2.3 s, 1 mm^3^ isotropic voxels. To perform the correction of magnetic field, the fieldmap scans (opposite phase encoded EPIs) were obtained. Task T2*-weighted scan was performed using Simultaneous multislice (SMS) imaging technique, characterized by 397 volumes, TE = 0.03 s, TR = 2 s, Slice Thickness = 2.5 mm, 60 slices acquired in interleaved order, Multiband acceleration factor = 2.

The MID task was programmed in the Presentation software.

### Behavioral data analysis

All analyses of behavioral data were performed using the Python programming language (version 3.9.6) and the following libraries: pandas (version 1.5.1), numpy (version 1.22.1), scipy (version 1.7.3), statsmodels (version 0.13.5), matplotlib (version 3.6.2) and seaborn (version 0.13.2). To assess the effect of distinctness and eveningness on psychometric parameters, the Ordinary Least Squares (OLS) regression analysis across multiple dependent variables was performed. The Variance Inflation Factor (VIF) for both predictors was ∼ 1, so well below the threshold of 5, dispelling multicollinearity concerns. The values were distributed normally according to the Kolmogorov-Smirnov test (*p* > 0.05). For each model homoscedasticity, the uniform variance of residuals across the range of predicted values, was confirmed via the Breusch-Pagan test and the Durbin-Watson statistic was used to check the autocorrelation among residuals and attest to the independence of errors. Together, these diagnostic tests confirmed the critical assumptions underlying the linear regression model, establishing a robust foundation for our analysis. The Ordinary Least Squares (OLS) regression model was fitted using the statsmodels library. Median split and two-way ANOVA were used to compare higher / lower-distinctness and higher / lower-eveningness individuals. Data normalization was performed using MinMaxScaler from scikit-learn library (version 1.2.2). Reaction times during the MID task were calculated using the Z-score for each data point. A Z-score greater than 3 or less than -3 was considered indicative of an outlier.

### MRI data analysis

Image processing and statistical analyses were done in the FMRIB Software Library (FSL^70^). Anatomical scans were reoriented to the standard (MNI152) orientation, bias-field corrected (RF/B1-inhomogeneity-correction), and non-linearly registered to the standard space, which was followed by brain-extraction, and segmentation^71^. In turn, the fieldmap files were prepared using FSLMERGE and TOPUP. The fMRI data were analyzed using the standard pipeline in the FEAT module (FMRI Expert Analysis Tool^72,73^). The first three volumes of the fMRI time series were discarded to allow for signal equilibration. The subsequent procedures included motion, slice timing, and distortion correction, removal of non-brain tissue, spatial smoothing with a Gaussian kernel of 8 mm FWHM and high-pass temporal filtering (90 s). Registration to the MNI152 standard space was obtained with a non-linear approach. The time-series were analysed with the general linear model (GLM), controlling for the temporal autocorrelation and motion during the scan. All explanatory variables (EV) were defined with a 3-column format and double-gamma HRF convolution.

Two types of analyses were performed. Primarily, we ran group-level whole-brain models using FMRIB’s Local Analysis of Mixed Effects (FLAME 1). Maps (z-stat) for each contrast were thresholded at *p* <.005 (*Z* = 2.8) with cluster correction *p* threshold set to 0.05. The biological clock data, i.e. eveningness and distinctness, were used as factors of interest.

In addition to the whole-brain models, we also performed region-of-interest (ROI) analyses for ventral tegmental area (VTA). The decision to use such an approach for the midbrain area was guided by its relatively small size, which deemed the cluster-extent significance thresholding a highly underpowered method. Therefore, VTA mask was created through thresholding a probabilistic ROI derived from an earlier work^74^ using the 0.999 cut-off, which was followed by its binarization and resampling to the resolution of fMRI data. The mask was used in Featquery module of FSL to extract the mean parameter estimates for all the contrasts of interest from the whole-brain analyses. The OLS linear model was run in Python statsmodel (version 0.13.5), using the VTA parameter estimates as the dependent variables, with eveningness and distinctness serving as their predictors.

## Data availability statement

We uploaded all data used in described analyses to the Openneuro database (https://openneuro.org/) and made it available in open-access – dataset entitled “Monetary Incentive Delay task - structural and functional images of 37 men; study of associations between circadian characteristics (eveningness, distinctness) and affective processing”^75^.

## Acknowledgments

We would like to express gratitude to Dawid Droździel, Marta Rodziewicz, and Bartosz Kossowski, PhD, from the Laboratory of Brain Imaging, Nencki Institute of Experimental Biology, Polish Academy of Sciences for their invaluable technical assistance with MRI data collection.

## Authors contributions

All authors conceived the project. All authors interpreted the results and revised the manuscript. **PS**: Conceptualization, Study design, Methodology, Participant recruitment, Data collection, Formal analysis, Writing - Original Draft, Visualization, Figures preparation, Funding acquisition, **MRZ**: Conceptualization, Methodology, Formal analysis, Writing - Original Draft, **JL**: Participant recruitment, Data collection, Writing - review & editing, **AES**: Writing - review & editing, Supervision, **PB**: Writing - review & editing, Supervision, **IS**: Conceptualization, Study design, Methodology, Writing - review & editing, Supervision.

## Supplementary data

**Supplementary Table S1.** The Ordinary Least Squares (OLS) regression results for distinctness and eveningness as predictor variables. Statistically significant correlations (at the p<0.05 level) are bolded and underlined. Psychometric parameters were assessed using the Sensitivity to Punishment and Sensitivity to Reward Questionnaire (SPSRQ), NEO-Five Factor Inventory (NEO-FFI), Generalized Anxiety Disorder (GAD-7), Positive and Negative Affect Schedule (PANAS). For details, see: Methods. Abbreviations: CI - confidence interval, D-W - Durbin-Watson statistics, B-P - Breusch-Pagan statistics.

|  | coef | Std err | t | P>\|t\| | 95% CI<br>[0.025; 0.975] |
| --- | --- | --- | --- | --- | --- |
| **Age** ($R^2 = 0.086$, $F = 1.595$, $p = 0.218$)<br>D-W = 2.694, B-P = 0.0975, B-P p-value = 0.952 | | | | | |
| Distinctness | 0.1861 | 0.112 | 1.666 | 0.105 | -0.041; 0.413 |
| Eveningness | -0.0518 | 0.074 | -0.704 | 0.487 | -0.202; 0.098 |
| **Sensitivity to punishment** ($R^2 = 0.228$, $F = 5.027$, $p = 0.0122$)<br>D-W = 1.801, B-P = 2.235, B-P p-value = 0.327 | | | | | |
| Distinctness | 0.4032 | 0.164 | 2.457 | **<u>0.019</u>** | 0.070; 0.737 |
| Eveningness | -0.2263 | 0.108 | -2.091 | **<u>0.044</u>** | -0.446; -0.006 |
| **Sensitivity to reward** ($R^2 = 0.078$, $F = 1.431$, $p = 0.253$)<br>D-W = 1.746, B-P = 3.107, B-P p-value = 0.211 | | | | | |
| Distinctness | 0.1449 | 0.091 | 1.595 | 0.120 | -0.040; 0.329 |
| Eveningness | 0.0304 | 0.060 | 0.507 | 0.615 | -0.091; 0.152 |
| **Neuroticism** ($R^2 = 0.096$, $F = 1.808$, $p = 0.0.179$)<br>D-W = 1.953, B-P = 0.290, B-P p-value = 0.865 | | | | | |
| Distinctness | 1.3615 | 0.379 | 3.590 | **<u>0.001</u>** | 0.591; 2.132 |
| Eveningness | 0.0963 | 0.250 | 0.385 | 0.703 | -0.412; 0.605 |
| **Extraversion** ($R^2 = 0.279$, $F = 6.578$, $p = 0.00385$)<br>D-W = 2.235, B-P = 1.469, B-P p-value = 0.480 | | | | | |
| Distinctness | 0.1530 | 0.398 | 0.385 | 0.703 | -0.656; 0.962 |
| Eveningness | 0.4849 | 0.262 | 1.847 | 0.073 | -0.048; 1.018 |
| **Openness** ($R^2 = 0.186$, $F = 3.875$, $p = 0.0305$)<br>D-W = 2.212, B-P = 0.966, B-P p-value = 0.617 | | | | | |
| Distinctness | 0.8870 | 0.337 | 2.634 | **<u>0.013</u>** | 0.203; 1.571 |
| Eveningness | 0.1793 | 0.222 | 0.807 | 0.425 | -0.272; 0.631 |
| **Agreeableness** ($R^2 = 0.049$, $F = 0.8793$, $p = 0.424$)<br>D-W = 2.121, B-P = 1.383, B-P p-value = 0.501 | | | | | |
| Distinctness | 0.5156 | 0.391 | 1.319 | 0.196 | -0.279; 1.310 |
| Eveningness | -0.0470 | 0.258 | -0.182 | 0.857 | -0.571; 0.477 |
| **Conscientiousness** ($R^2 = 0.057$, $F = 1.025$, $p = 0.370$)<br>D-W = 1.890, B-P = 8.944, B-P p-value = 0.011 | | | | | |
| Distinctness | -0.4100 | 0.407 | -1.008 | 0.321 | -1.237; 0.417 |
| Eveningness | -0.2630 | 0.268 | -0.980 | 0.334 | -0.808; 0.282 |
| **GAD** ($R^2 = 0.021$, $F = 0.3649$, $p = 0.697$)<br>D-W = 1.862, B-P = 1.389, B-P p-value = 0.499 | | | | | |
| Distinctness | 0.1478 | 0.182 | 0.810 | 0.423 | -0.223; 0.518 |
| Eveningness | -0.0360 | 0.120 | -0.299 | 0.767 | -0.280; 0.208 |
| **Positive affect** ($R^2 = 0.013$, $F = 0.2155$, $p = 0.807$)<br>D-W = 2.528, B-P = 0.207, B-P p-value = 0.902 | | | | | |
| Distinctness | -0.0126 | 0.376 | -0.033 | 0.974 | -0.777; 0.752 |
| Eveningness | -0.1623 | 0.248 | -0.654 | 0.517 | -0.667; 0.342 |
| **Negative affect** ($R^2 = 0.088$, $F = 1.645$, $p = 0.208$)<br>D-W = 1.923, B-P = 2.533, B-P p-value = 0.282 | | | | | |
| Distinctness | 0.5375 | 0.298 | 1.804 | 0.080 | -0.068; 1.143 |
| Eveningness | -0.0505 | 0.197 | -0.257 | 0.799 | -0.450; 0.349 |
| **BAS Drive** ($R^2 = 0.014$, $F = 0.2368$, $p = 0.790$)<br>D-W = 2.388, B-P = 2.255, B-P p-value = 0.324 | | | | | |
| Distinctness | -0.0990 | 0.150 | -0.660 | 0.513 | -0.404; 0.206 |
| Eveningness | 0.0215 | 0.099 | 0.217 | 0.829 | -0.180; 0.223 |
| **BAS Fun seeking** ($R^2 = 0.081$, $F = 1.493$, $p = 0.239$)<br>D-W = 1.753, B-P = 7.596, B-P p-value = 0.022 | | | | | |
| Distinctness | 0.0081 | 0.091 | 0.090 | 0.929 | -0.176; 0.192 |
| Eveningness | 0.1030 | 0.060 | 1.721 | 0.094 | -0.019; 0.225 |
| **BAS Reward** ($R^2 = 0.066$, $F = 1.199$, $p = 0.314$)<br>D-W = 2.511, B-P = 1.959, B-P p-value = 0.376 | | | | | |
| Distinctness | 0.1335 | 0.139 | 0.962 | 0.343 | -0.149; 0.416 |
| Eveningness | -0.1142 | 0.092 | -1.247 | 0.221 | -0.300; 0.072 |
| **BIS** ($R^2 = 0.297$, $F = 7.191$, $p = 0.00249$)<br>D-W = 1.462, B-P = 1.509, B-P p-value = 0.470 | | | | | |
| Distinctness | 0.6283 | 0.195 | 3.220 | **<u>0.003</u>** | 0.232; 1.025 |
| Eveningness | -0.2724 | 0.129 | -2.117 | **<u>0.042</u>** | -0.534; -0.011 |
| **Attention deficits** ($R^2 = 0.238$, $F = 5.296$, $p = 0.00995$)<br>D-W = 2.083, B-P = 2.727, B-P p-value = 0.256 | | | | | |
| Distinctness | 1.3589 | 0.513 | 2.648 | **<u>0.012</u>** | 0.316; 2.402 |
| Eveningness | 0.6078 | 0.338 | 1.796 | 0.081 | -0.080; 1.296 |

**Supplementary Table S2.** Two-way ANOVA results for pairs of positive-negative psychometric traits. Groups of higher / lower distinctness and higher / lower eveningness were obtained using a median split. Statistically significant associations (at the p<0.05 level) are bolded and underlined.

|  | Source of Variation | SS | df | F | PR(>F) |
| --- | --- | --- | --- | --- | --- |
| **Negative affect** | **Distinctness** | 125.485 | 1.0 | 4.266 | **<u>0.047</u>** |
|  | **Eveningness** | 18.873 | 1.0 | 0.642 | 0.429 |
|  | **Interaction** | 3.316 | 1.0 | 0.113 | 0.739 |
|  | **Residual** | 970.759 | 33.0 |  |  |
| **Positive affect** | **Distinctness** | 20.078 | 1.0 | 0.412 | 0.526 |
|  | **Eveningness** | 3.813 | 1.0 | 0.0782 | 0.782 |
|  | **Interaction** | 16.423 | 1.0 | 0.337 | 0.566 |
|  | **Residual** | 1609.059 | 33.0 |  |  |
| **Neuroticism** | **Distinctness** | 445.032 | 1.0 | 8.713 | **<u>0.006</u>** |
|  | **Eveningness** | 0.423 | 1.0 | 0.008 | 0.928 |
|  | **Interaction** | 163.603 | 1.0 | 3.203 | 0.083 |
|  | **Residual** | 1685.611 | 33.0 |  |  |
| **Extraversion** | **Distinctness** | 11.880 | 1.0 | 0.217 | 0.645 |
|  | **Eveningness** | 182.109 | 1.0 | 3.321 | 0.078 |
|  | **Interaction** | 13.513 | 1.0 | 0.246 | 0.623 |
|  | **Residual** | 1809.559 | 33.0 |  |  |
| **BIS** | **Distinctness** | 93.595 | 1.0 | 6.163 | **<u>0.018</u>** |
|  | **Eveningness** | 18.487 | 1.0 | 1.217 | 0.278 |
|  | **Interaction** | 7.896 | 1.0 | 0.520 | 0.476 |
|  | **Residual** | 501.184 | 33.0 |  |  |
| **BAS** | **Distinctness** | 16.212 | 1.0 | 0.605 | 0.442 |
|  | **Eveningness** | 0.00054 | 1.0 | 0.0 | 0.9964 |
|  | **Interaction** | 30.367 | 1.0 | 1.133 | 0.295 |
|  | **Residual** | 884.66 | 33.0 |  |  |

**Supplementary Table S3.** Results of the re-conducted neuroimaging analyses – excluding 3 participants, who obtained Athens Insomnia Scale values higher than 10, which indicated potential chronic sleep deprivation. Abbreviations: B, bilateral. ^a^Whole-brain analysis corrected for multiple comparisons with cluster-level family-wise error rate approach. ^b^Region-of-interest analysis.

| Contrast | MNI (x, y, z) | Area | Voxels | Statistics | p-value |
| --- | --- | --- | --- | --- | --- |
| Loss anticipation > implicit baseline | 0, 24, 62 | B superior frontal gyrus<br>B supplementary motor area | 268 | Z = 3.42 | 0.029<sup>a</sup> |
| Loss anticipation > win anticipation | 0, -17, -14 | B ventral tegmental area | 32 <sup>a</sup> | T = 2.899 | 0.007<sup>b</sup> |

**Supplementary Table S4.** Two-way ANOVA results for behavioral measurements during Monetary Incentive Delay task. Groups of higher / lower distinctness and higher / lower eveningness were obtained using a median split. Median reaction times for each trial type (all trials, big win trials, small win trials, big loss trials, small loss trials) and total score were treated as dependent variables. We found no statistically significant relationships.

|  | Source of Variation | SS | df | F | PR(>F) |
| --- | --- | --- | --- | --- | --- |
| Reaction time – all trials | Distinctness | 525.229 | 1.0 | 0.738 | 0.397 |
|  | Eveningness | 2413.277 | 1.0 | 3.391 | 0.075 |
|  | Interaction | 135.969 | 1.0 | 0.191 | 0.665 |
|  | Residual | 23488.545 | 33.0 |  |  |
| Reaction time – big win trials (+ 5 PLN) | Distinctness | 388.547 | 1.0 | 0.418 | 0.522 |
|  | Eveningness | 1868.169 | 1.0 | 2.012 | 0.165 |
|  | Interaction | 0.793 | 1.0 | 0.0009 | 0.977 |
|  | Residual | 30642.662 | 33.0 |  |  |
| Reaction time – small win trials (+ 1 PLN) | Distinctness | 865.821 | 1.0 | 0.971 | 0.332 |
|  | Eveningness | 3123.983 | 1.0 | 3.504 | 0.070 |
|  | Interaction | 8.155 | 1.0 | 0.009 | 0.924 |
|  | Residual | 29421.504 | 33.0 |  |  |
| Reaction time – big loss trials (- 5 PLN) | Distinctness | 743.600 | 1.0 | 1.279 | 0.266 |
|  | Eveningness | 1326.027 | 1.0 | 2.282 | 0.140 |
|  | Interaction | 721.849 | 1.0 | 1.242 | 0.273 |
|  | Residual | 19179.768 | 33.0 |  |  |
| Reaction time – small loss trials (- 1 PLN) | Distinctness | 772.739 | 1.0 | 0.661 | 0.422 |
|  | Eveningness | 1799.279 | 1.0 | 1.539 | 0.224 |
|  | Interaction | 403.019 | 1.0 | 0.345 | 0.561 |
|  | Residual | 38578.458 | 33.0 |  |  |
| Total score | Distinctness | 11.032 | 1.0 | 0.026 | 0.874 |
|  | **Eveningness** | 10.813 | 1.0 | 0.025 | 0.875 |
|  | **Interaction** | 306.624 | 1.0 | 0.711 | 0.405 |
|  | **Residual** | 14239.332 | 33.0 |  |  |

